# Notch and LIM-homeodomain protein Arrowhead regulate each other in a feedback mechanism to play a role in wing and neuronal development in *Drosophila*

**DOI:** 10.1101/2024.09.16.613220

**Authors:** Jyoti Singh, Dipti Verma, Bappi Sarkar, Maimuna Sali Paul, Mousumi Mutsuddi, Ashim Mukherjee

## Abstract

Notch pathway is an evolutionarily conserved signaling system that operates to influence an astonishing array of cell fate decisions in different developmental contexts. To identify novel effectors of Notch signaling, we analyzed the whole transcriptome of *Drosophila* wing and eye imaginal discs in which an activated form of Notch was overexpressed. A LIM homeodomain protein Arrowhead (Awh) was identified as a novel candidate which plays a crucial role in Notch mediated developmental events. *Awh* alleles show strong genetic interaction with Notch pathway components. Awh loss-of-function upregulates Notch targets Cut and Wingless. Awh gain-of-function downregulates Notch targets by reducing the expression of ligand, Delta. Consequently, the expression of Wingless effector molecule Armadillo and its downstream targets, Senseless and Vestigial, also gets downregulated. Awh overexpression leads to ectopic expression of *engrailed*, a segment polarity gene in the anterior region of wing disc, leading to patterning defects. Additionally, Notch gain-of-function mediated neuronal defects get significantly rescued with Awh overexpression. Activated Notch inhibits Awh activity, suggesting a regulatory loop between Awh and Notch. Additionally, the defects caused by Awh gain-of-function were remarkably rescued by Chip, a LIM interaction domain containing transcriptional co-factor. The present study highlights the novel feedback regulation between Awh and Notch.

## Introduction

The Notch pathway is an evolutionarily conserved signaling system and it is highly pleiotropic in nature because it regulates a spectrum of cellular events such as cell fate determination, differentiation, proliferation, apoptosis and stem cell maintenance during development (Andersson et al., 2011; Artavanis-Tsakonas et al., 1999; Kopan & Ilagan, 2009). Notch is synthesized as a 300 kDa precursor protein that is subjected to its first cleavage during its maturation in the trans-Golgi network by furin-like convertases (S1 cleavage), resulting in a N-terminal extracellular domain (NECD) of 180 kDa and a C-terminal transmembrane intracellular domain (NTM) of 120 kDa (Blaumueller et al., 1997). This heterodimeric Notch receptor is transported to the plasma membrane where it interacts with ligands of the DSL family (*Drosophila* Delta and Serrate (Jagged in mammals) and *C. elegans* LAG-2) expressed in the adjacent cell. This receptor-ligand binding leads to second proteolytic cleavage (S2) by a disintegrin and metalloprotease (ADAM) family of metalloproteases in the extracellular region of the NTM (Brou et al, 2000). This is followed by an intramembranous cleavage (S3) by γ-secretase complex (Presenilin, Nicastrin, PEN-2, and APH-1) and results in the release of Notch intracellular domain (NICD) from the membrane (De Strooper et al., 1999; Struhl & Greenwald, 1999). Then, Importin-α3 mediates the translocation of NICD to the nucleus (Sachan et al., 2013). NICD associates with the DNA binding protein CSL (mammalian CBF1/ *Drosophila* Su(H)/ *C. elegans* Lag-1) in the nucleus and facilitates the displacement of transcriptional co-repressors. The NICD-CSL complex then recruits Mastermind and other transcriptional co-activators leading to activation of Notch target genes such as the *Enhancer of Split [E(spl)]* complex genes in *Drosophila* (Bray & Bernard, 2010; Fortini & Artavanis-Tsakonas, 1994; Kopan & Ilagan, 2009; Wu et al., 2000). These bHLH transcription factors, inturn, repress *achaete-scute complex* (As-C) proneural genes. Depending on the cellular context, NICD-CSL activation complex can also activate other Notch target genes such as *wg, cut, string/CDC-25, c-myc,* etc. (Bray & Bernard, 2010; Krejcí et al., 2009). The increasingly complex regulatory mechanisms of Notch signaling account for the multitude of functions exhibited by Notch during development.

Similarly, Wingless (Wg) acts as a ligand and activates Wg signaling by binding to the receptor Frizzled. Wg signaling is an evolutionarily conserved pathway that regulates pivotal roles in cell fate determination during embryonic development. Upon binding with the receptor, it leads to dual phosphorylation of Lrp6 by GSK-3-β (Shaggy) and casein kinase-1 (CKI) in cytosol and shifting of the whole complex to plasma membrane along with Dishevelled (Dsh). This complex aids in stabilization of β-catenin (Armadillo). β-catenin moves to the nucleus and leads to activation of downstream target genes *senseless (sens), distal-less (dll) and vestigial* (*vg*). Inthe absence of Wg, a complex of Axin, APC (Adenomatous polyposis coli), GSK3-β, CKI andβ-catenin is present in cytosol. Armadillo is phosphorylated by GSK3-β in this complex and directed to proteasomal degradation (Komiya & Habas, 2008).

These two signaling pathways, Notch and Wg, are the major regulators of compartmentalisation and fate determination of wing blade, dorsal-ventral boundary, wing hinge region as well as wing vein development of *Drosophila melanogaster* (Couso et al., 1994). During early developmental events *engrailed (en)*, a homeodomain containing segment polarity gene is expressed in cells adjacent to wg expressing cells and maintains wg transcription. Wg is also crucial to maintain En levels, indicating an autoregulation of its expression through paracrine feedback loop (Heemskerk et al., 1991). Feedback regulation is a fundamental approach that helps cells to uphold proper homeostasis and govern a range of cellular processes. It is a dynamic system that provides adaptability, resilience and stability to cellular system.

Here, we present functional characterization of a novel effector of Notch signaling identified through transcriptome analysis. At this end, whole transcriptome profiling of the *Drosophila* wing and eye imaginal discs overexpressing activated form of Notch was carried out (Paul et al., 2018). Arrowhead (Awh), a LIM homeodomain (LIM-HD) protein, was identified as a novel candidate which plays a crucial role in Notch mediated developmental events. LIM-HD proteins are characterized by presence of two tandem LIM domains [named for the first defining members: Lin-11, Isl-1, and Mec-3], and a homeodomain (Freyd et al., 1990; Karlsson et al., 1990; Gehring et al., 1994). The LIM domain consists of two unique zinc fingers that are rich in cysteine and connected by a short two-amino acid linker. It acts as a transcription factor. It is a highly conserved molecule across the taxa; its mammalian ortholog, lhx8 has been identified to have a role in neuronal development via WNK signaling pathway (Sato & Shibuya, 2013). Awh is well known for its role in establishment of proper number of precursor cells in incorporate imaginal tissues such as abdominal histoblast and salivary gland imaginal rings in *Drosophila*. Interestingly, both loss-of-function and gain-of-function alleles of Awh affect early developmental events. The ectopic expression of Awh in excorporate imaginal discs leads to elimination of cells (Curtiss & Heilig, 1997).

Here we report that a LIM homeodomain protein Arrowhead (Awh) was identified through a transcriptome analysis from the activated Notch-induced wing and eye imaginal discs in which Awh was downregulated. *Awh* mutant alleles displayed strong genetic interaction with *Notch* pathway components. It was seen that Awh loss-of-function upregulates Notch targets, Cut and Wg. Gain-of-function of Awh downregulates these Notch targets, Cut and Wg, and reduction of these targets was mediated through the downregulation of the ligand, Delta without altering Notch receptor levels. Here we also show that Awh plays a major role in wing development by regulating Notch-mediated Wingless signaling. Additionally, neuronal defects caused due to Notch gain-of-function were get rescued with Awh overexpression. Interestingly, activated Notch inhibits Awh activity, highlighting the novel feedback regulation between Awh and Notch.

## RESULTS

### *Awh* genetically interacts with Notch pathway components

To decipher the functional implications of Awh in the Notch pathway, first we checked whether mutations in *Awh* and Notch pathway components display genetic interaction in trans-heterozygous combinations. We used *Awh16, Awh63Ea-1,* and *Awh63Ea-E12* functional null alleles generated by using Ethyl Methane Sulfonate mediated base pair substitution (Curtiss & Heilig, 1995, 1997; Wohlwill & Bonner, 1991). Notch null allele *N54l9* showed significantly penetrant wing notching phenotype, bringing *Awh* alleles in this background in trans-heterozygous condition resulted in rescue of wing nicking, indicating enhancement of Notch function (Fig. 1.A1-A4). Mild notching phenotype of male flies of Notch hypomorphic allele *Nnd3* got rescued with *Awh* alleles in trans-heterozygous condition (Fig. 1.B1-B4). Driving the dominant negative form of Mastermind in the C96 domain led to enhanced notching; the phenotype got rescued by reducing the level of Awh in trans-heterozygous condition (Fig. 1.C1-C4). *C96-GAL4* driven expression of dominant negative form of *Notch* showed fully penetrant severe wing notching. This wing nicking phenotype of Dominant negative form of Notch was rescued to a great extentin trans-heterozygous combination with *Awh* alleles (Fig. 1.D1-D4). The present study provides evidence of significant genetic interaction between *Awh* and components of *Notch* pathway.

**Fig. 1.**
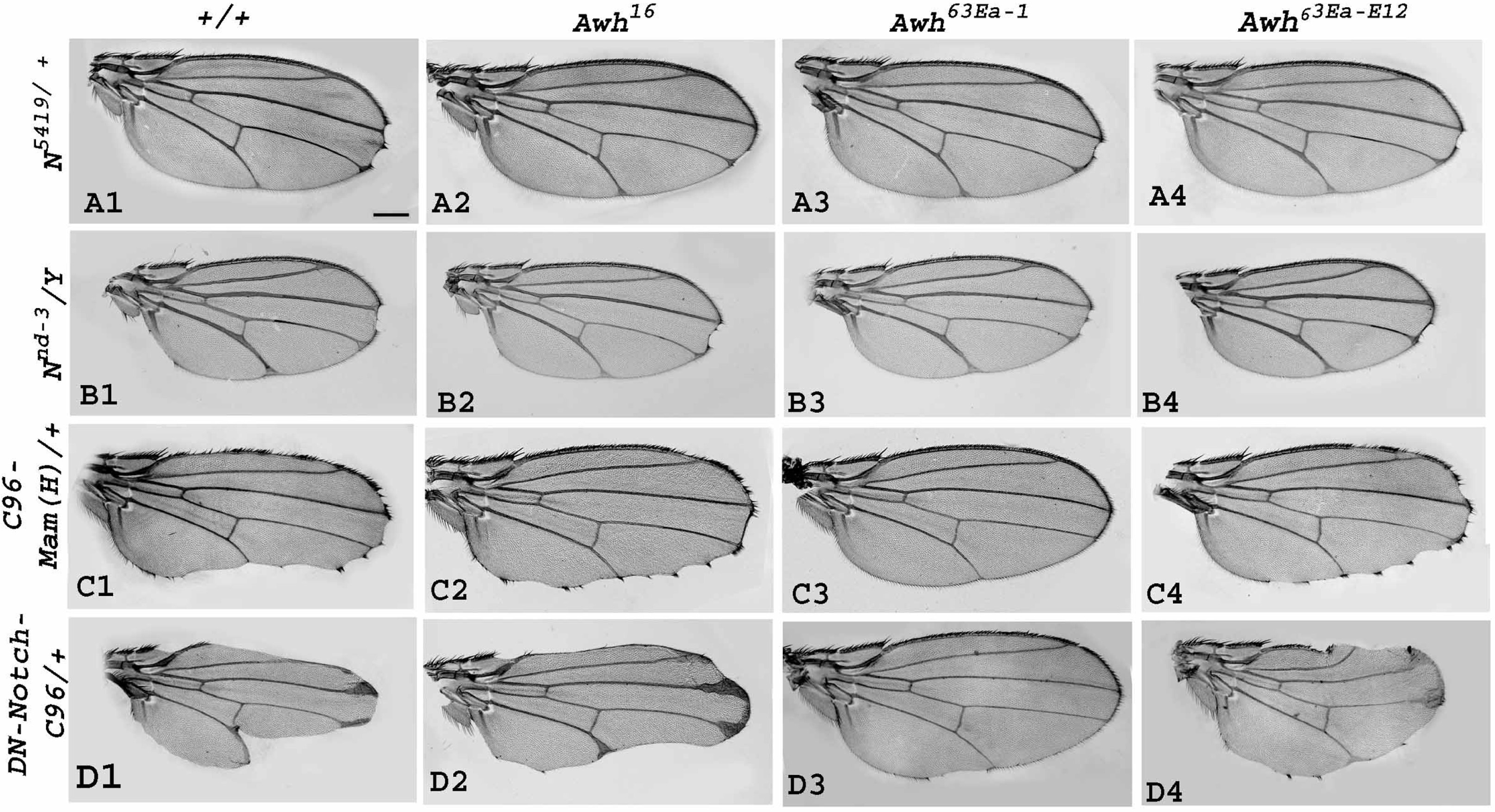
Genetic interactions of *Awh* alleles with Notch pathway components. Representative wings with genotypes are shown. Wings with *N54l9* heterozygotes showed the wing nicking phenotype, trans-heterozygous combination of *N54l9* with different *Awh* alleles *Awh16, Awh63Ea-1, Awh63Ea-E12* showed reduced wing notching (A1-A4). Notch hypomorph allele *Nnd3* showed mild notching at wing margin, whereas in trans-heterozygous combinations with *Awh* alleles, it showed rescue in wing phenotype (B1-B4). *C96-GAL4* driven Dominant-negative Mastermind (*MamH*) in heterozygous condition displayed wing notching phenotype, which was reduced in trans-heterozygous combinations with *Awh* alleles; (C1-C4). Overexpression ofdominant negative form of Notch in DV boundary exhibit severe wing notching, which was rescued in combination with *Awh* alleles (D1-D4).). Scale Bar: 150µm (A1-D4)

### Loss-of-function of Awh upregulates Notch Signaling

To decipher the loss-of-function effects of Awh, we checked the status of Notch signaling activity by investigating the expression levels of downstream target of Notch, Wg in Awh loss-of-function background. Notch induces Cut and Wg expression at the dorsoventral (DV) boundary of the developing wing disc (Neumann & Cohen, 1996). Downregulation of Awh in dorsal region using *ap-GAL4* leads to a slight increase in Wg levels as shown by immunostaining (Fig. 2.A1-A4).

**Fig. 2.**
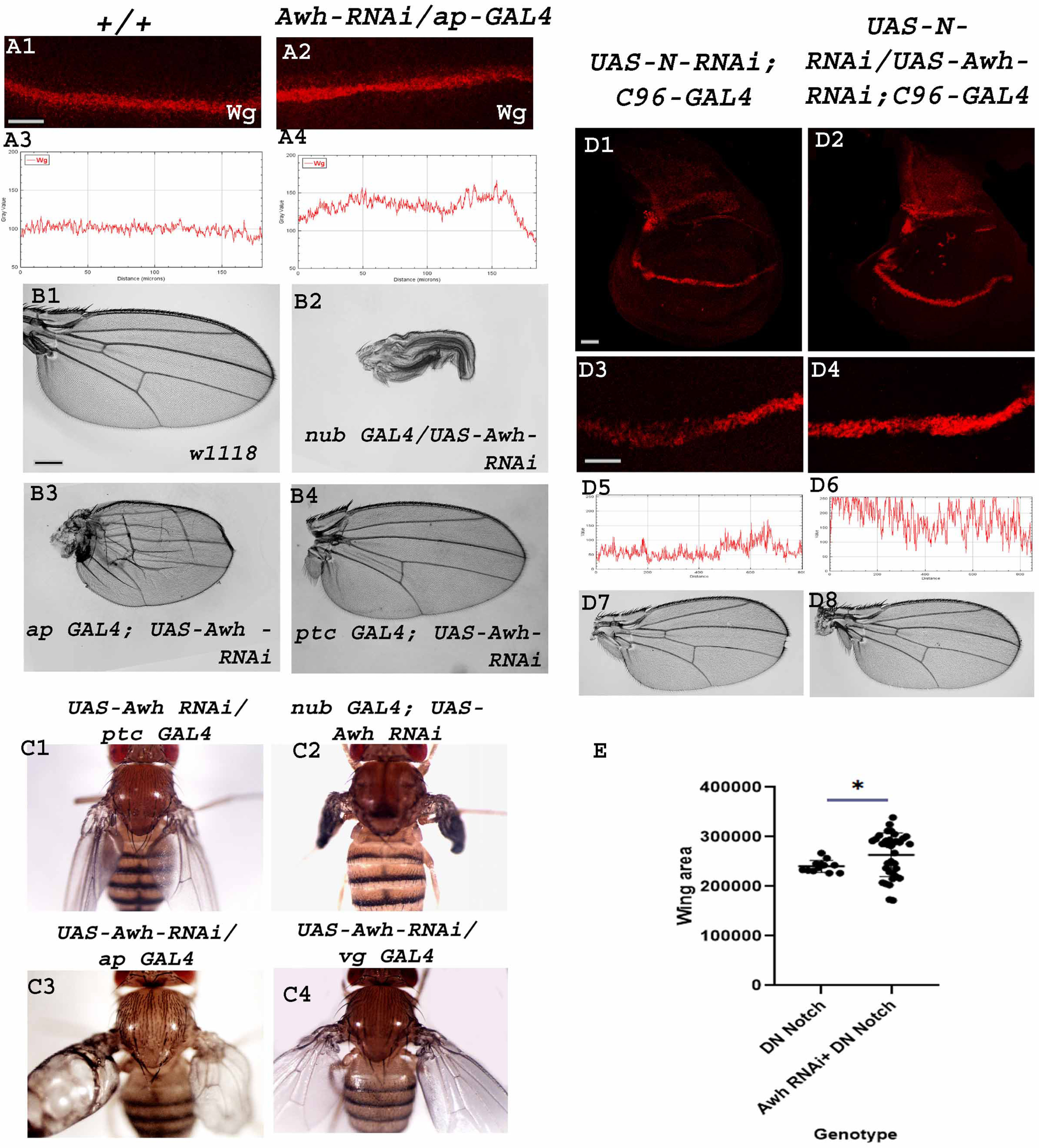
Downregulation of Awh leads to increased Notch activity: Downregulation of Awh in dorsal region using *ap-GAL4* transgene led to increased expression of Wg as compared to wild type discs (A1-A2). RGB plot profiles indicating the intensity of Wg in A1 and A2, respectively (A3-A4). Adult wing images shows effect of Awh loss-of-function on wing development. Awh downregulation leads to paralysed wings (*nub-GAL4*), blister formation (*ap-GAL4*), defects in vein formation, loss of Anterior cross vein (*ap-GAL4* and *ptc-GAL4*) and erect wing phenotype (*ptc-GAL4, nub-GAL4, ap-GAL4, vg-GAL4*) (B1-C4). Downregulation of Notch RNAi using *C96-GAL4* leads to downregulation of Cut (D1, high magnification-D3) and notching in adult wing (D7). Bringing Awh-RNAi in the background has rescued the Cut expression (D2, High magnification-D4) and wing notching phenotype. RGB profile plot shows the intensity of Cut (D5, D6). Graph shows the rescue in *DN-Notch::C96 GAL4* wing area upon bringing *UAS-Awh* in background (E). Scale Bar: 30 μm (A1-A2, D1-D4); 150 μm (B1-B4, D7-D8)

Downregulation of Awh in the pouch region leads to formation of paralysed wing where the wings fails to open up. Its loss-of-function in dorsal region using *ap-GAL4* transgene leads to blister formation, severe defects in wing vein formation, extra vein material at hinge region anda severe crumpling. Similarly, Awh downregulation in AP boundary leads to increased vein material between 3rd-4th vein, and loss of ACV (Anterior cross vein). It is interesting to notice that Awh downregulation in AP boundary, wing pouch, dorsal and ventral region of wings leads to erect wing phenotype (Fig. 2.B1-C4).

Further we checked the effect of Awh downregulation in Notch downregulation background. Downregulation of Notch in DV boundary using RNAi line leads to decreased expression of Cut in wing imaginal disc and notching at wing margin in adult (28% flies shows notching, no. of flies observed =80). Interestingly, upon bringing Awh loss-of-function in the background the Notch loss-of-function the Cut expression was rescued and no notching was observed in adults (no. of flies observed=80). (Fig. 2.D1-D8) Similar experiment was performed using Dominant negative form of Notch. It was observed that there was a rescue in wing size upon bringing Awh-RNAi in *UAS-DN-Notch; C96-GAL4* background (Fig. 2.E). From the dataset it is evident that Awh loss-of-function has significantly rescued Notch loss-of-function phenotype.

### Awh downregulates Notch Signaling

To explore the involvement of Awh in Notch pathway we checked the status of Notch signaling activity by investigating the expression levels of downstream targets of Notch, Cut and Wg in Awh overexpressed wing imaginal discs. Overexpression of Awh by *ptc-GAL4* in anterior-posterior (AP) boundary showed complete reduction of Notch targets, Cut and Wg at the AP and DV intersection in comparison to wild type control as shown in images (Fig. 3.A1-B4) (no.of discs examined= 30). RGB profile plots display the decreased fluorescence intensity of Cut and Wg in the regions with overexpressed Awh compared to the internal control (Fig. 3.C1, C2).In addition, we checked the transcript levels of Notch targets, Enhancer of Split [*E(spl*)] complex genes in Awh overexpressed condition using Real-time PCR. *E(spl*) contains seven transcription units *m8*, *m7*, *m5*, *m3*, *m*β, *m*γ, and *m*δ that encode basic helix-loop-helix containing transcription factors (Bailey & Posakony, 1995). The expression of all seven E(spl)-HLH transcripts were downregulated in Awh overexpression background in comparison to *Gmr-GAL4/+* control (Fig. 3.D).

**Fig. 3.**
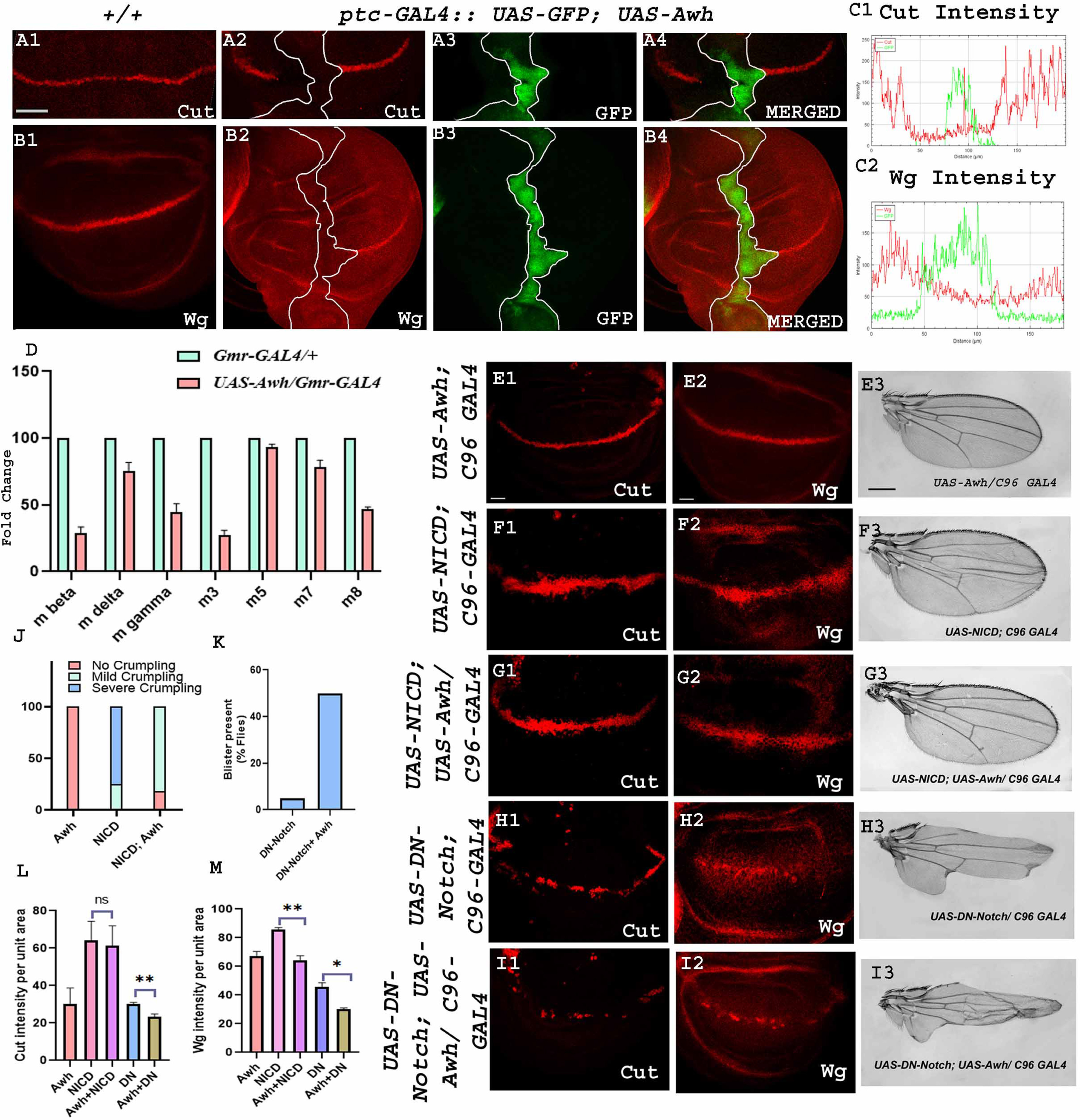
Downregulation of Notch signaling by Awh overexpression: Overexpression of Awh in AP boundary using *ptc-GAL4::UAS-GFP* transgene led to a significant reduction in level of Cut and Wingless as compared to wild type wing discs, GFP marks the Awh overexpression domain, Merged images show Cut and Wg expression along with GFP, White line marks the boundary of Awh overexpression (A1-B4). RGB plot profiles indicating the intensity of A4, and B4, respectively; Red marks the Cut and Wg intensity and green marks the Awh overexpression domain (C1, C2). Level of Notch target, *E(spl)* complex gene transcripts (*m*β*, m*γ*, m*Δ, *m3, m5, m7, m8*) are shown in Awh overexpression background, *GAL4* control was used. Reduction in all seven *E(spl)* complex genes transcripts was observed in Awh overexpression condition (D). Cut and Wg expression is shown in C96 driven Awh overexpression condition (E1, E2). NICD overexpression with *C96-GAL4* showed increased Cut and Wg expression in wing discs (F1, F2) and crumpled wings in adult (F3), Co-expressionof Awh and NICD rescue the Cut and Wg expression to some extent (G1, G2), rescue in wing crumpling was also observed (G3). Graph showing the ratio of mild and severe crumpling in wings with different combination of Awh and NICD (J). Overexpression of Dominant negative Notch in DV boundary showed significant reduction in level of Notch targets Cut and Wg (H1, H2), adult with overexpressed Dominant negative Notch showed severe notching (H3), Overexpressing Awh together with Dominant negative form of Notch led to significant disruption in targets Cut and Wg (I1, I2) as well as in adult phenotype (I3) as compared to Notch-DN, The adult wing also displays increase in notching, and presence of blisters (H1-I3).Graph showing the increase in percentage of flies with blister (K). Intensity profiling of Cut and Wg was shown in different combination of Awh, NICD and DN-Notch driven with *C96-GAL4* (L, M). Scale Bar: 30 μm (A1-B4, E1-E2, F1-F2, G1-G2, H1-H2, I1-I2); 150µm (E3,F3,G3,H3,I3).

**Fig. 4.**
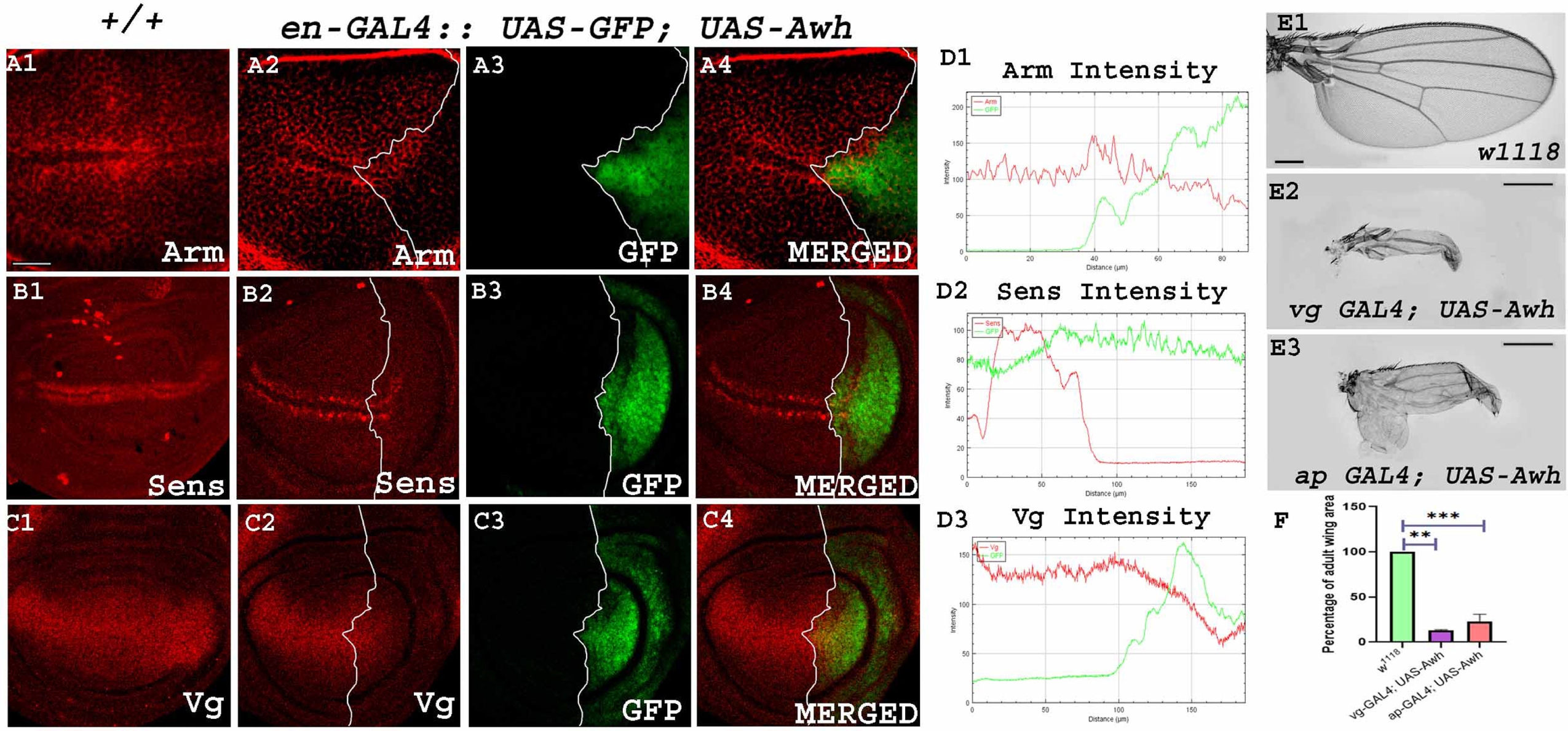
Awh modulates the expression of Wg signaling components: Representative wing disc shows Arm expression in wild type control discs (A1), Awh overexpression in the posteriordomain of wing disc using *en-GAL4* led to significant disruption in Arm expression, GFP marks the posterior domain of the third instar wing disc, merged image shows the expression of Arm and GFP in Awh overexpression condition, White line marks the boundary of Awh overexpression (A2-A4). Representative wing disc shows Sens expression in wild type control discs (B1), Awh overexpression led to decrease in Sens expression in posterior compartment, GFP marks the posterior domain of the third instar wing disc, merged image shows the expression of Sens and GFP in Awh over-expression background. White line marks the boundary of Awh overexpression (B2-B4). Representative wing disc shows Vg expression in wild type discs (C1), Awh overexpression in posterior compartment downregulated Vg expression, GFP marks the posterior domain of the third instar wing disc, Merged image shows expression of Vg and GFP in Awh over-expression, White line marks the boundary of Awh overexpression (C2-C4). RGB plot showing intensity of Arm, Sens and Vg in Awh overexpression, Red marks the intensity of Arm, Sens and Vg and green marks the Awh overexpression domain (D1-D3). Representative wings with genotypes are shown. Awh overexpression in ventral and dorsal regions by *vg-GAL4* and *ap-GAL4* (E2, E3) led todrastically small adult wings in comparison to wild type wings (E1). Graph showing the comparison of Adult wing area in control and Awh overexpression wings (F). Scale Bar: 30μm(A1-C4); 3cm (E1-E3).

Further, we checked the loss and gain-of-function effects of Notch on the Awh overexpressed wing imaginal discs. As modulation in Notch levels using *ptc-GAL4* leads to embryonic lethality, *C96-GAL4,* a comparatively weak GAL4 transgene was used for interaction studies. Awh overexpression in the DV boundary using *C96 GAL4* resulted in mild defects in bristle patterning while no significant changes were observed in Cut and Wg expression (Fig. 3.E1-E3). Overexpression of NICD alone in DV boundary led to increased expression of Notch targets, Cut and Wg level hence consequently led to mild crumpling and irregular bristle patterning in adult wing whereas co-expressed *UAS-Awh* along with *UAS-NICD* in DV boundary showed decrease in wing size and crumpling of adult wing as shown in graph (no. of wings examined = 80) (Fig. 3.J). The levels of Notch targets Cut and Wg were also rescued in the co-expression background (no. of discs examined = 30) (Fig. 3.F1-G3). Further we checked the interaction between Awh and Notch Dominant Negative. Dominant Negative form of Notch is the Notch receptor lacking Notch activation domain (NICD) (Rebay et al., 1993). Overexpressing Notch dominant negative in *C96-GAL4* domain led to severe wing notching. Co-expression of Awh and dominant negative form of Notch further worsens the wing phenotype, the severity of notching increases, presence of blisters was also observed in 50% of flies as shown in graph (no. of wings examined = 80) (Fig. 3.K). This prompted to check the level of Notch targets in this background and we noticed that the levels of Cut and Wg were significantly downregulated in *UAS-DN-Notch; UAS-Awh/ C96-GAL4* in comparison with *UAS-DN-Notch; C96-GAL4* (no. of discs examined = 30) (Fig. 2.I1-I3). This result showed that Awh downregulates Notch signaling. Quantification of Notch targets, Cut and Wg was done inthe same background. About 3 discs were used for quantification using ImageJ software where integrated density/area of the domain was used for quantification purpose. Unpaired t-test was performed to determine the significance of our finding. Intensity profiling further validated the role of Awh in downregulation of Notch signaling (Fig. 3.L, M).

### Awh overexpression modulates Wingless signaling

Since Awh overexpression led to the downregulation of Notch target Wg, we checked the effect of Awh overexpression on Wg signaling activity and wing development. Wg plays a central role in pattern formation during development (Couso et al., 1994). In wild type tissue when Wg binds to receptor Frizzled it leads to the stabilization of Arm protein, this stabilized Arm enters the nucleus and activates downstream target genes: *sens, dll, and vg* (Couso et al., 1994; Mosimann et al., 2009; Neumann & Cohen, 1997). In the absence of ligand Wg, the multiprotein destruction complex: APC, Axin, and GSK-3 degrades Wg effector, Arm by phosphorylation (Mosimann et al., 2009). Interestingly, overexpression of Awh in posterior compartment of third instar wing disc showed a strong disruption in the level of Arm that consequently led to the downregulation of short range and long range targets Sens and Vg in comparison to the wild type discs (no. of discs examined = 30) (Fig. 4.A1-C4). RGB plot showsa significant downregulation of Wg targets in the domain of Awh overexpression (Fig. 4.D1-D3). Further, Awh gain-of-function mediated changes in adult wing was observed.*vg-GAL4* and *ap-GAL4* driven Awh overexpression in ventral and dorsal region of *Drosophila* wing discs, respectively, forms vestigial wing phenotype reminiscent of wingless and vestigial loss-of-function phenotype that further strengthen up our study (no. of wings examined = 80) (Fig. 4.E1-E3) (Simmonds et al., 1997). Quantitative analysis of adult wing size was performed using ImageJ software and graph was plotted using Graph Pad Prism 8 software. Statistical analysis highlights the significant reduction in wing size in Awh overexpression background (Fig. 4.F).

We also checked whether the influence of Awh on Wg signaling is Notch-mediated. To this end, activated form of Notch was provided in Awh gain-of-function background and the expression of Wg was observed. Temperature sensitive *ptc-GAL4::UAS-GFP* was used for our study. As *ptc-GAL4* mediated NICD overexpression is embryonic lethal temperature sensitive GAL80 system was employed for temporal regulation of UAS transgene. As expected Awh gain-of-function completely abolished Wg expression in the AP domain. Overexpression of Notch intracellular domain led to increased expression of Wg along the AP boundary. When *UAS-Notch-ICD* and *UAS-Awh* were coexpressed, Wg expression was significantly reduced in wing discs compared to that of only NICD overexpressed wing imaginal discs. This result indicated that Awh regulated Wg signaling might be controlled through Notch pathway(Supplementary Figure. S1).

### Awh regulates wing patterning

Awh overexpression in the AP boundary of wing discs using *ptc-GAL4* driver leads to several types of patterning defects in wing discs. 70% of wings shows drastic inward folding of wing imaginal discs, while 15% discs manifest small duplication and 15% shows complete duplication in the wing pouch (no. of discs examined = 60) (Fig. 5.A2-A3). We checked the expression of Cut and Wg in the duplicated region. The duplicated part showed ectopic Cut and Wg expression (no. of discs examined = 8) (Fig. 5.B1, C1 High magnification in B2, C2). Clonal experiments were performed to interrogate the spatial regulation of Awh gain-of-function mediated wing duplication, we found that the clone specifically formed in the dorsal compartment of wing disc showed ectopic expression of Cut and Wg. The clones of the ventral region showed no change in expression of these Notch targets, while clones formed on the DV boundary region showed loss of Cut and Wg (no. of discs examined = 20) (Supplementary Figure. S2). Earlier studies have shown that the expression pattern of Awh transcripts in stage 14-16 of embryo is very much similar to segment polarity genes (Curtiss & Heilig, 1997). Segment polarity genes, *wg* and *engrailed* (*en*) play profound roles in wing development (Baker, 1988; Couso et al., 1994). In this context, along with Wg expression we also checked En expression and found En is also ectopically expressed in duplicated wing pouch area (no. of discs examined = 8) (Fig. 5.D1, High magnification in D2) in Awh overexpressed wing imaginal discs. This kind of duplication and ectopic expression of Wg and En clearly indicates the critical role of Awh in wing development.

**Fig. 5.**
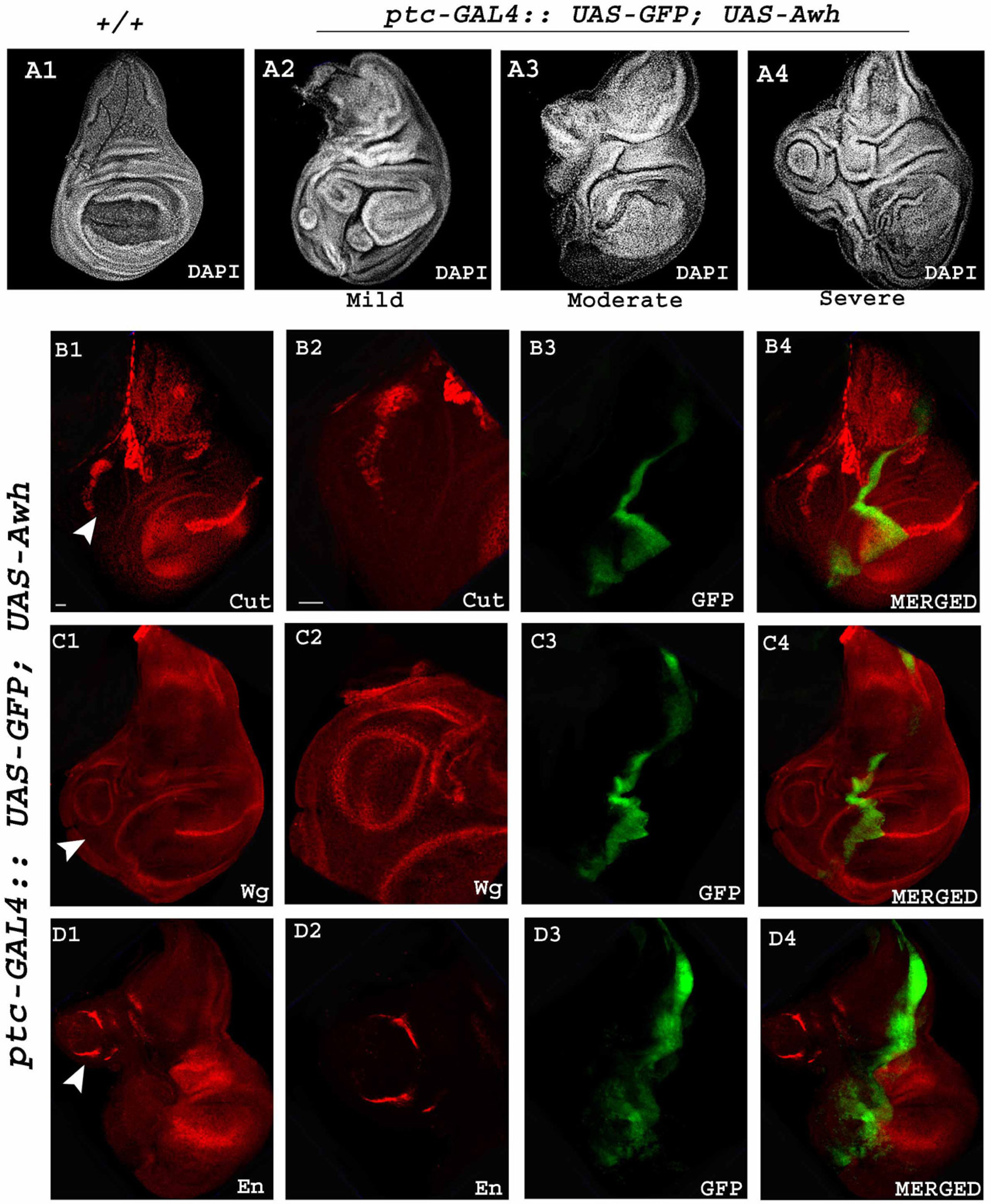
Awh regulates wing patterning. Awh overexpression in anterior posterior boundary using *ptc-GAL4::UAS-GFP* leads to drastic defects in wing discs morphology in comparison to control discs (A1) as shown by DAPI stain, defects were categorised as mild, moderate, and severe (A2-A4). Representative wing disc showing ectopic expression of Cut, Wg, and En in the anterior compartment of third instar wing disc in Awh overexpression condition (B1,C1,D1, High magnification B2,C2,D2). GFP marks the domain of Awh overexpression (B3,C3,D3), Merged images show expression of Cut and GFP, Wg and GFP, En and GFP (B4, C4, D4). Scale Bar: 30 μm (A1-D4).

### Right amount of Chip expression is crucial for Awh function

It has been shown earlier that Awh prevents retinal differentiation and promotes differentiation to ventral head tissue in the regions of eye disc and Awh needs transcriptional co-factor Chip for this function (Roignant et al., 2010). We have shown earlier that Chip interacts with Notch and it plays a major role in Notch-induced DV margin formation and cell proliferation (Sachan et al., 2015). These observations prompted us to check the role of Chip in Awh-mediated wing development.

*vg-GAL4* driven Awh overexpression in the ventral domain of the wing discs displayed severe loss of wing tissue, whereas the co-expression of Awh and Chip in the ventral domain of wing discs rescued this wing phenotype. Similarly, Awh overexpression in the pouch region using *nub-GAL4* led to pupal lethality whereas the co-expression of Awh and Chip led to rescue from pupal lethality (no. of wings examined = 50) (Fig. 6.A1-A6) which showed that the right amount of Chip is important for proper Awh function. Graphical representation of the percentage of wing area showed significant recovery of adult wing size in Awh-Chip co-expression (Fig. 6.A7-A8). We also observed recovery in wing disc size and morphology in Awh-Chip co-expression background (Supplimentary Figure S3). Similarly we found that Awh overexpression in the posterior region of the wing imaginal discs by *en-GAL4* led to abolition of Notch targets Cut and Wg, while the co-expression of Awh and Chip in posterior domain restored the Cut and Wg (no. of discs examined = 25) (Fig. 6.B1-B12) levels. RGB profile plots highlights the similar observations (Fig. 6.D1-D6). Thus, it is evident again that a critical amount of Chip is important for Awh function.

**Fig. 6.**
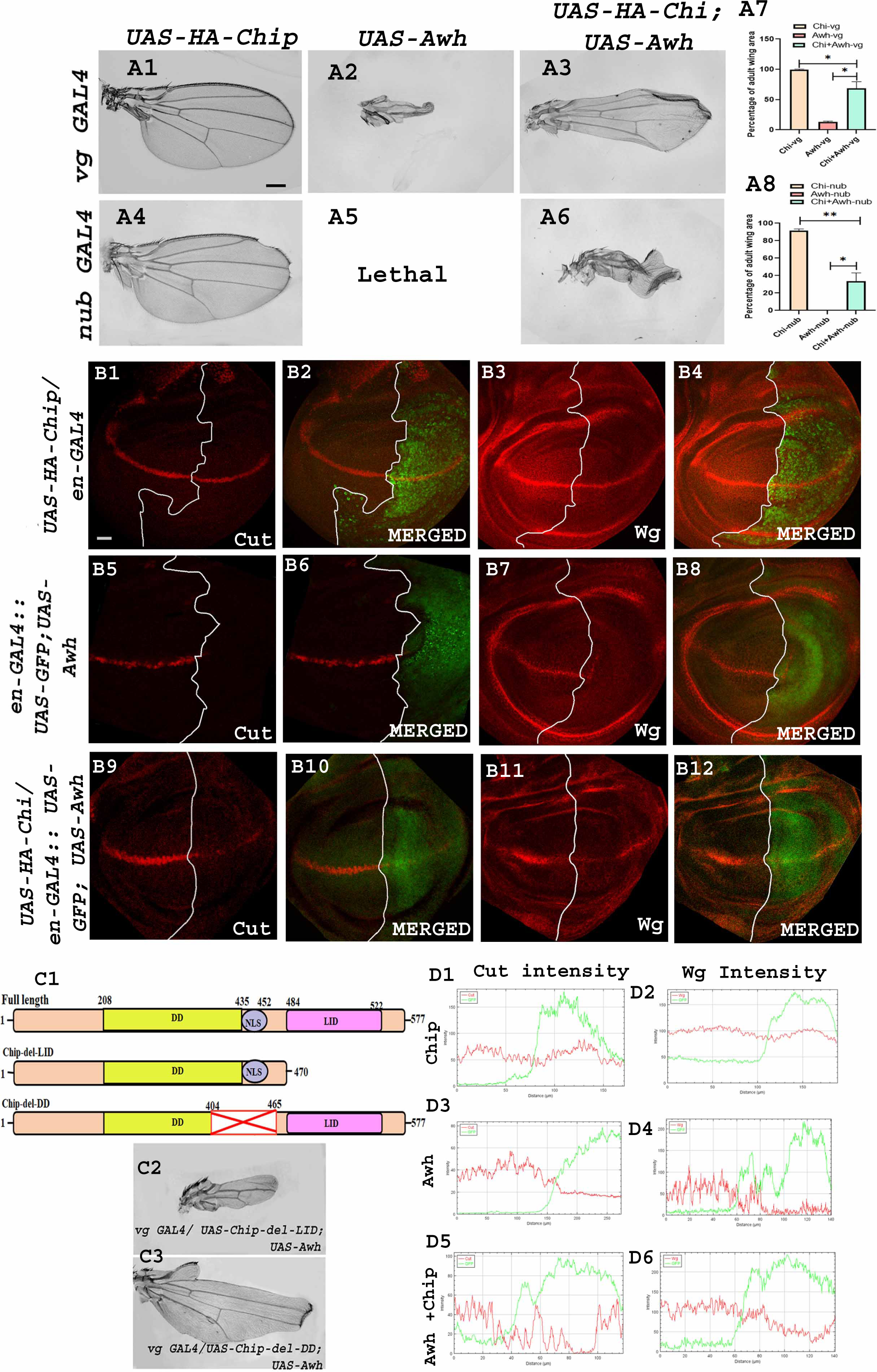
Chip rescue Awh gain-of-function phenotype. Representative wings with genotypes are shown. Chip overexpression in the ventral region of the wing showed no significant change (A1), Awh overexpression in the ventral region showed severe loss of wing tissue (A2) while co-expression of Chip and Awh rescued the wing phenotype (A3). Chip overexpression in pouch region shows mild notching (A4), Awh overexpression using *nub-GAL4* led to pupal lethality (A5) while co-expression of Awh and Chip in pouch region rescue the lethality, the adult wing showed severe crumpling (A6). Graph showing the rescue of wing area in Awh and Chip co-expression (A7-A8). Representative wing disc showing expression of Cut and Wg in HA-Chip overexpression in posterior compartment (B1, B3). White lines shows the boundary of Chip overexpression (B1-B4). Expression of Cut and Wg was completely lost in Awh gain-of-function condition (B5, B7). White lines shows the boundary of Awh overexpression (B5-B8). Co-expression of Chip and Awh restores the expression of Cut and Wg to some extent (B9, B11). White lines shows the boundary of Awh and Chip co-expression (B9-B12). RGB profile plot shows intensity of Cut and Wg in different genetic combinations, Red marks the Cut, Wg intensity and green marks the domain of overexpression (D1-D6). Domain structure of full length Chip and deletion lines are shown (C1). Overexpressing Chip with deleted LIM domain along with Awh showed no recovery in wing size, Overexpressing Chip with deleted dimerisation domain along with Awh showed recovery in wing size (C2-C3). Scale Bar: 150 μm (A1-A6, C2-C3); 30 μm (B1-B12).

**Fig. 7.**
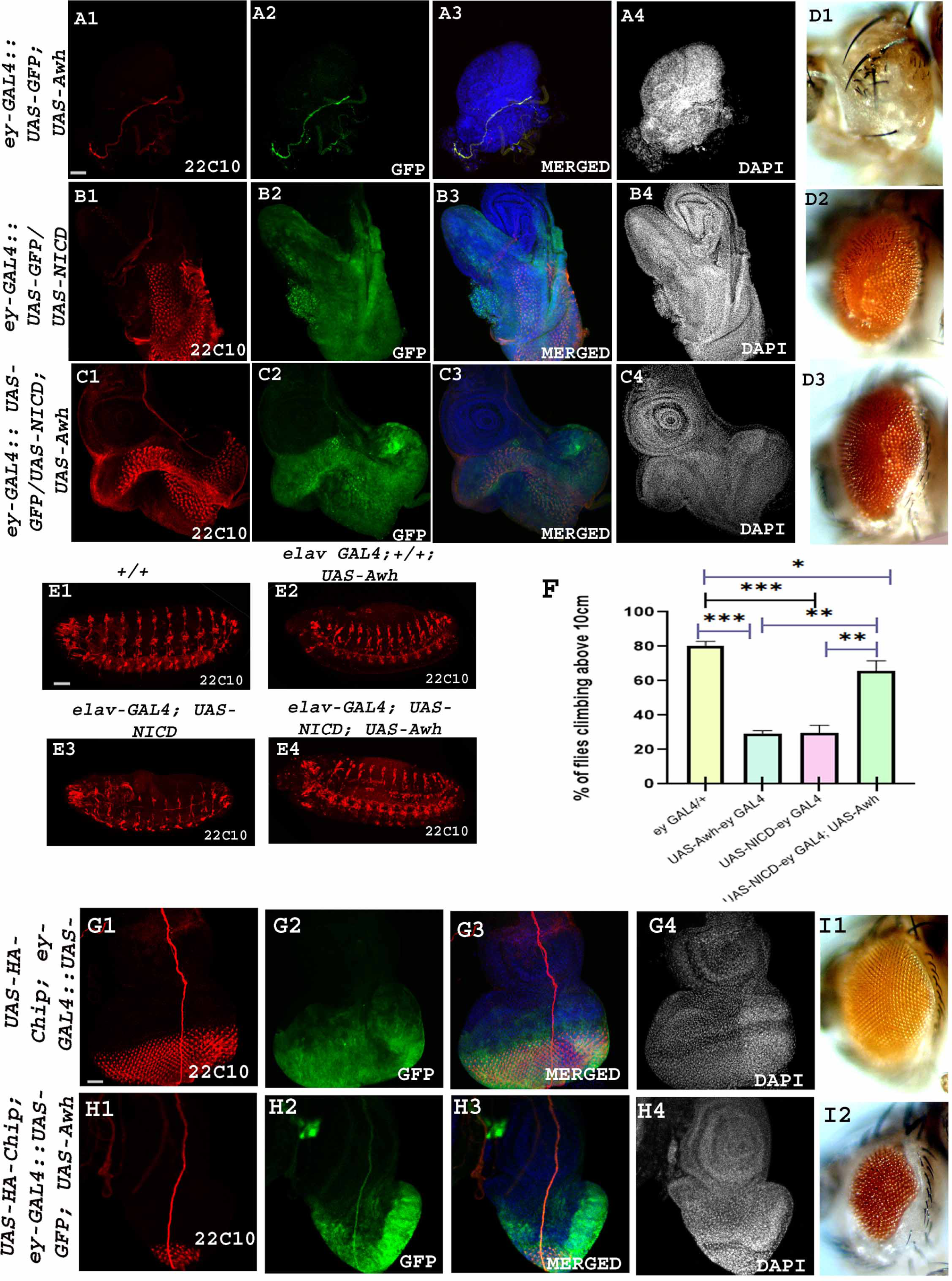
Awh regulates Notch-mediated neuronal defects. Overexpression of Awh using *ey-GAL4* led to loss of neuronal tissue marked with 22C10 (A1), GFP marks the domain of Awh overexpression (A2), Merged image shows 22C10, GFP and DAPI (blue) together, DAPI shows eye disc morphology (A3, A4), Overexpression of NICD using *ey-GAL4* led to disrupted 22C10 expression(B1), GFP marks the domain of NICD overexpression (B2), Merged image shows 22C10, GFP and DAPI (blue) together, DAPI showed hyperproliferative eye disc (B3 B4). Co-expression of Awh and NICD using *ey-GAL4* led to restoration of 22C10 expression (C1), GFP marks the co-expression domain (C2), Merged image shows 22C10, GFP and DAPI (blue) together, DAPI shows recovery of eye morphology (C3, C4). Representative adult eye showing ablated eye in Awh overexpression condition (D1), NICD overexpressed adult shows hyperproliferative eye (D2), while the adult eye shows rescue in Awh and NICD co-expressed background (D3). Similar results were observed in embryos using the 22C10 antibody. Overexpression of NICD using *elav-GAL4* showed significant defects in the ventral nerve cord of stage 14 embryo in comparison to wild type embryo. Co-expression of Awh and NICD led to significant rescue of ventral nerve cord (E1-E4). Climbing ability was monitored using climbing assay, only Awh and only NICD overexpressed flies showed defect in climbing ability, while co-expression of Awh and NICD showed significant rescue in the climbing defects (F). Co-expression of Chip and Awh using *ey-GAL4* rescued the Awh overexpression mediated photoreceptor cell defects shown by 22C10 (H1), GFP marks the co-expression domain (H2),Merged image shows 22C10, GFP and DAPI (blue) together, DAPI shows recovery of eye morphology (H3,H4). Scale Bar: 30 μm (A1-C4, E1-E4, G1-H4).

Chip contains a self-dimerisation domain and a LIM binding domain. In order to check which domain of Chip is crucial for interaction with Awh, different Chip deletion lines were co-expressed with Awh. Two deletion lines of Chip were used, one line has deleted the LIM binding domain while the other one has deleted the dimerization domain. Different domains of full-length Chip protein and the deletion lines were shown in Fig. 6.C1. Bringing Awh in combination with Chip deleted LIM binding domain led to no significant rescue while its co-expression with Chip deleted dimerization domain led to a remarkable rescue in adult wing phenotype (no. of wings examined = 80) (Fig. 6.C2, C3). This dataset suggests to us that the LIM binding domain of Chip is crucial for Awh functioning.

### Awh regulates Notch-mediated Neuronal defects

Earlier the role of Awh in neuronal development was shown. Awh overexpression using *Da-GAL4* leads to defects in the Futsch expression pattern (Sato & Shibuya, 2013). Futsch is a microtubule binding protein, helps in formation of synaptic buttons at neuromuscular junctions (Roos et al., 2000). Notch being identified as a neurogenic gene plays a crucial role in neurogenesis and neuron maintenance (Poulson, 1937). To study the functional application of Awh-Notch regulation we checked the level of neuronal marker, Futsch/22C10 in different combinatorial manner. Increasing the level of Awh in the eye discs using *ey-GAL4* caused significant loss of neuronal cells as well as loss of eye tissue in adults. On the contrary, NICD overexpression in the eyeless region led to disruption in the Futsch pattern and proliferative eye phenotype in adults. The combinatorial effect of Awh and activated Notch in the eye led to betterment of eye phenotype, and the pattern of neuronal marker 22C10 also got restored (Fig. 7.A1-D3). Similar results were obtained in *Drosophila* embryos stage 14 using *elav-GAL4* (Fig. 7.E1-E4). To further validate the study climbing ability of different sets of flies were observed.The only Awh and activated Notch flies showed defects in their climbing ability while bringing them together restored their climbing behaviour in comparison to GAL4 control flies (no. of flies examined = 50) (Fig. 7.F).

Further, we explored the effect of Chip on Awh mediated neuronal defects. Over-expression of Chip in eyeless domain shows normal pattern of photoreceptor cells. Co-expressing Awh with Chip rescues Awh mediated loss of photoreceptor cells (no. of disc examined = 25). A prominent rescue is also observed in adult eye (Fig. 7.G1-I2).

### Awh regulates Notch Signaling by affecting Delta expression

Since overexpression of Awh led to reduced expression of Notch targets, we wanted to dissect out its effect on the status of Notch receptor. We checked the level of endogenous Notch protein in the wing discs in which Awh was overexpressed in the posterior compartment using *en-GAL4* transgene. Notch expression and localization remained unaltered in the Awh overexpressing domain of the wing disc (Fig. 8.A1-A4). This shows that Awh downregulates Notch signaling without affecting Notch receptor levels.

As the presence and binding of DSL ligands is crucial for Notch receptor activation (Kopan & Ilagan, 2009) and Delta is already shown to regulate Awh transcripts (Curtiss & Heilig, 1997), it prompted us to check the level of Delta in Awh overexpressed discs. The expression pattern of Delta was analysed while overexpressing Awh in the posterior compartment of the third instar wing disc. Awh gain-of-function illustrates a remarkable decrease in Delta in the posterior compartment (Fig.8.B1-B4). Further genetic interaction studies were performed to fortify our observation. *Delta* alleles showed vein thickening phenotype. The severity of vein thickening was further enhanced upon bringing *Awh* alleles in background confirming the genetic interaction between the molecules (Fig.8.C1-C4). Wings were categorised into mild, moderate and severe with different degree of vein thickening. Scoring of the adult wing was done. *DlBX6* shows wings with 45% mild and 55% moderate vein thickening. Bringing Awh allele *Awh16, Awh63Ee-1, Awh63Ea-12* led to mild vein thickening to 37%, 4%, 0% moderate vein thickening to 63%, 45%, and 9.5% and severe vein thickening in 0%, 50%, 90.5% of wings respectively (Fig. 8.E) (no. of wings observed = 50). This study suggests that Awh mediated Notch signaling downregulation is Delta-mediated.

To strengthen up our hypothesis we analysed the level of Dl in Awh loss-of-function condition. Awh was downregulated in the entire wing pouch using *nub-GAL4* and checked the expression of Delta. It was observed that there was an increase in Dl level (Fig. 8.F1). Similarly,downregulation of Awh in dorsal compartment leads to a significant disruption in Dl expression(Fig. 8.G1). We also checked the level of Dl transcripts upon downregulating Awh in eye imaginal disc using *Gmr-GAL4* transgene. An increase in Delta transcripts was observed in Awh downregulation condition (Fig. 8.H). This dataset confirms that there is a negative feedback loop between Awh and Delta.

### Awh activity is regulated by activated Notch

Awh is reported to cause programmed cell death and loss of adult structures in overexpressed condition (Curtiss & Heilig, 1997). We also observed increased Acridine Orange positive cells which marked the dying cells when Awh was overexpressed in different domains of wing discs (Supplementary Figure. S4). It was seen that Awh mediated cell death is caspase dependent (Supplimentary Figure. S5). Inhibition of Caspase mediated cell death shows recovery in wingsize but no significant rescue was observed in DV boundary confirming that Awh gain-of-function mediated Notch regulation is independent of Cell death (Supplimentary Figure. S6). *vg-GAL4* driven overexpression of Awh transgene in the ventral region shows increased expression of dying cells marked by DCP-1. Earlier it has been shown, overexpression of *NICD*in the ventral region shows a hyper-proliferative phenotype with no cell death in the wing disc(Ho et al., 2015). We have also seen the similar phenotype when NICD was overexpressed. Further, we have observed that if Awh is overexpressed along with Notch ICD, Awh overexpression induced cell death was also absent (no. of disc examined = 35)(Fig. 9.A1-C3). Awh transcripts level was shown to be significantly downregulated in activated Notch condition(Paul et al., 2018). Thus, it is prompting to speculate that feedback regulation between Awh and Notch may exist.

**Fig. 8.**
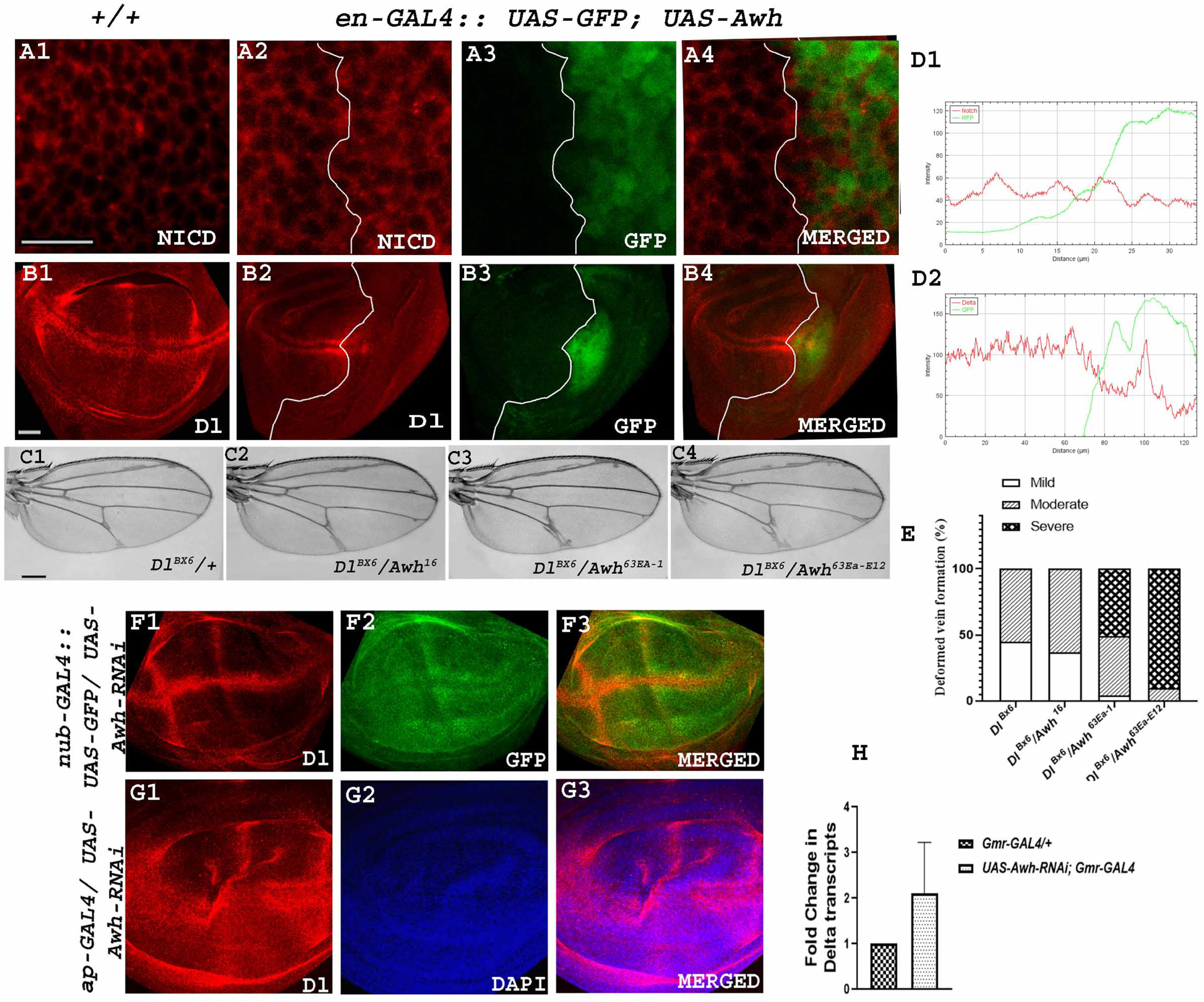
Awh affects Notch pathway by regulating Delta expression. Representative wing disc showing Notch receptor expression in wild type wing discs (A1). Awh overexpression in the posterior domain of wing disc using *en-GAL4:: UAS-GFP* led to no change in expression of Notch receptor, GFP marks the posterior domain of the third instar wing disc, Merged imageshows the expression of Notch and GFP in Awh over-expression, White line marks the boundary of Awh overexpression (A2-A4). Representative wing disc showing Delta expressionin wild type wing discs (B1). Awh over-expression in the posterior domain of wing disc using*en-GAL4* led to significant reduction in Dl expression, GFP marks the posterior domain of thethird instar wing disc, Merged image shows the expression of Dl and GFP in Awh over-expression, White line marks the boundary of Awh overexpression (B2-B4). RGB profile plot demonstrate the intensity of Notch and Dl (marked with red) and Awh over-expression domain(marked with green) (D1-D2). Representative wings with genotypes are shown. *DlBX6* showed vein thickening phenotype, the thickening was significantly enhanced upon bringing Awh functional null *Awh16, Awh63Ea-1, Awh63Ea-12*in the background (C1-C4). Graph shows the percentage of flies with mild, moderate and severe level of vein thickening in different combination (E). Awh downregulation in pouch region using *nub-GAL4:: UAS-GFP* shows increased Delta expression (F1), GFP marks the domain (F2), Merged image shows the expression of Delta and domain of Awh downregulation. Delta expression in Awh loss-of-function using *ap-GAL4* is shown (G1), DAPI marks the disc morphology (G2) Merged imagesshows expression of Delta and DAPI (G3). Graph shows increase in Delta transcripts in Awh loss-of-function background (H). Scale Bar: 30 μm (A1-B4, F1-G3); 150 μm (C1-C4).

**Fig. 9.**
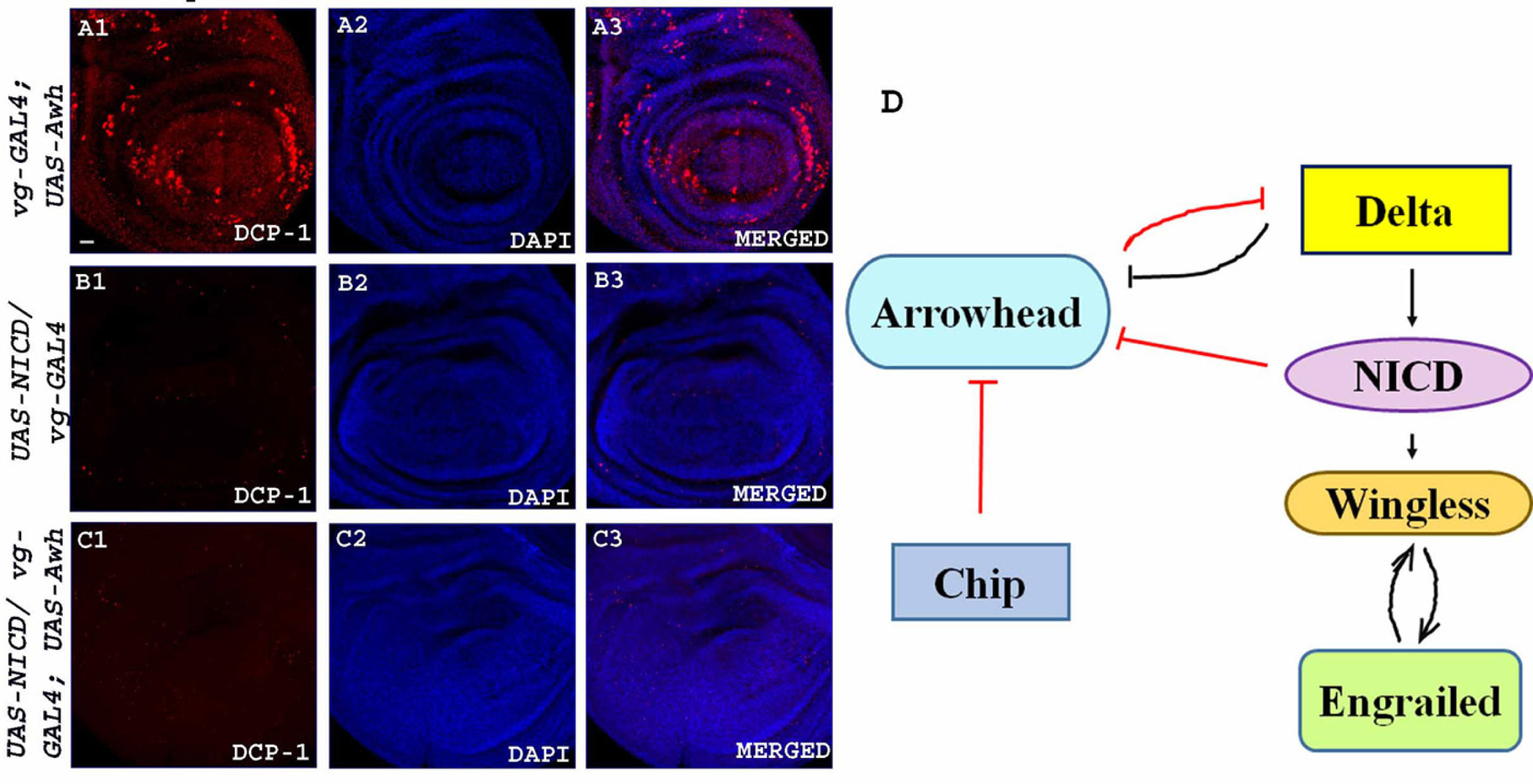
Awh activity is inhibited by activated Notch. Representative wing disc showing expression of dcp1-1 in Awh overexpression in the ventral region, morphology of disc was visualised with DAPI (A1-A3). NICD over-expression in ventral region showed no expression of dcp-1, DAPI shows the morphology of the disc (B1-B3) Awh and NICD co-expression alsoshowed no expression of dcp-1, DAPI shows the morphology of the disc (C1-C3). The model shows the summary of present work; the red arrows mark the novel finding, the black arrows are the established interactions (D). Scale Bar: 30μm (A1-C3).

## DISCUSSION

The development of multicellular organisms rely on precisely regulated multiple cellular signal transduction pathways. The disruption in any signaling cascade may lead to catastrophic events for cells. To escape from such deleterious events the cell has evolved multiple levels of intricate regulation of signaling pathways. Cross-talk between signaling pathways, signal integration and feedback mechanisms fine-tune the outcome of signaling pathways.

Here we present a strong genetic interaction between *Awh* and *Notch* alleles. Awh loss-of-functions upregulates Notch Signaling. Awh downregulates Notch Signaling without altering the Notch receptor levels. It downregulates Notch signaling activity by downregulating the ligand, Delta. As a consequence of Notch signaling downregulation, Awh overexpression also downregulates Notch target Wg. Earlier it was also shown that there is increased transcripts of Awh in *Delta* mutant embryos which supports the presence of a feedback loop between Notch and Awh (Curtiss & Heilig, 1997). Additionally, we found that the optimum amount of transcriptional co-factor Chip is critical for Awh function in wing patterning. Awh also rescues the neuronal anomalies caused by Notch gain-of-function. Further we observed that Awh activity gets compromised in activated Notch condition revealing a feedback mechanism between LIM-homeodomain transcription factor Awh and Notch.

Notch pathway has a profound role in different fundamental developmental events broadly classified as lateral inhibition, cell lineage decisions and boundary formation. Notch and Wg signaling pathways are crucial for DV boundary formation in both developing eye and wing imaginal discs. LIM homeodomain protein Apterous expression in the early wing primordium leads to expression of the Notch ligand Serrate in dorsal cells and restricts expression of another Notch ligand, Delta to ventral cells (Diaz-Benjumea & Cohen, 1995). Notch is symmetrically activated in cells on both sides of the DV compartment boundary by dorsally expressed Serrate and ventrally expressed Delta (de Celis et al., 1996; Diaz-Benjumea & Cohen, 1995; Doherty et al., 1996). Expression of the glycosyltransferase Fringe makes dorsal cells more sensitive to Delta and less sensitive to Serrate (Fleming et al., 1997; Moloney et al., 2000). Consequently, activated Notch induces Wg expression in cells along the DV boundary. Wg further activates expression of Serrate and Delta in nearby dorsal and ventral cells, respectively, and Serrate and Delta signal back to activate Notch and thereby maintain Cut and Wg expression along the DV boundary (Milán & Cohen, 2000, 2003). Thus, Notch and its ligands Delta and Serrate expression establishes a positive feedback loop that maintains signaling at the DV boundary in the wing imaginal discs. Similarly, a major role of Notch has been identified in boundary formation between the prospective somites dur ing vertebrate somitogenesis (Wahi et al., 2016) and at the midbrain– hindbrain boundary for organizer gene expression in chick embryos (Tossell et al., 2011). LIM-homeodomain protein Awh along with transcriptional co-factor Chip together plays a major role in restricting the eye field so that it helps in boundary formation between eye field and head surface at the ventral side (Roignant et al., 2010). Notch, Awh and Chip are very well conserved and their orthologues are present from *Drosophila* to Humans. Thus, we speculate that these proteins together fine-tune the boundary formation during metazoan development.

Notch was considered as a neurogenic gene since *Notch* mutants display a hyperplasia of the nervous system at expense of epidermal fate (Poulson, 1945). The orthologue of Awh, lhx6 has emerged as a probable candidate for Tourette syndrome, a neuro-muscular disease (Paschou et al., 2012). This prompted us to study the effect of Awh-Notch interplay on neuronal development. The neuronal defects of Notch gain-of-function were significantly reversed upon bringing Awh in the background. The climbing assay also supports the similar result. This observation opens up new avenues to study the Awh and Notch interaction in different neurodegenerative diseases in which dysregulation of the Notch pathway is involved.

The present study highlights the Awh and Notch feedback regulation and its functional implication in wing patterning, morphogenesis and neurogenesis.

## MATERIALS AND METHODS

### Drosophila genetics

All fly stocks were maintained on standard cornmeal/yeast/molasses/agar medium at 25 °C. *w1118* was used as wild-type controls. *UAS-Awh-RNAi* was a kind gift from Prof. L.S. Shashidhara, *C96GAL4::UAS-Mam(H),UAS-NICD, UAS-Notch DN* (Dominant Negative Notch) and Notch pathway components *Nnd-3, N54l9,* were kindly provided by Prof. S. Artavanis-Tsakonas, Department of Cell Biology, Harvard Medical School, *UAS-Awh* was obtained as a gift from Prof. Ella Preger-Ben Noon, Howard Hughes Medical Institute., Ashburn, VA, USA. *Awh63Ea-12* was a kind gift from Prof. Hiroshi Shibuya, Simon Fraser University,Canada. *Dl BX6*was requested from Prof. Alexey Veraksa, University of Massachusetts Boston, USA.*UAS-HA-Chip* was previously generated in the lab (Sachan et al., 2015). GAL4 driver lines: *ap-GAL4/CyO, C96-GAL4, en-GAL4:: UAS-GFP, ptc-GAL4::UAS-GFP, ptc-GAL4::UAS-GFP::GAL80ts, nub-GAL4, Gmr-GAL4, ey-GAL4::UAS-GFP, elav-GAL4, vg-GAL4, Awh16, Awh63Ea-1,UAS-Chip-*Δ*LID,* and *UAS-Chip-*Δ*DD* were procured from Bloomington *Drosophila* stock center. *vg-GAL4*::*UAS-GFP* was made with the help of appropriate genetic crosses. *nub GAL4::UAS-GFP* was requested from Cytogenetics Laboratory, Department of Zoology, BHU. All the crosses were performed at 25 °C.

To generate Awh gain-of-function clones, females of *hsp70-flp; Act FRT y FRT-GAL4* were crossed to *UAS-Awh* males. Heat shock was given at 37 °C for 12 minutes at 24 hrs after egg laying (AEL) and the third instar larval discs were analyzed for GFP-marked clones.

For studying overexpression of genes using temperature sensitive GAL80 system the desired crosses were allowed for egg laying at 18°C. When embryos mature into late first instar larvaethey were transferred to 24°C for 24hrs and again moved to 18°C. The larvae and adult offspring were further used for studies.

### Adult eye imaging and wing mounting

Flies were anaesthetized and placed on a bridged-slide. The eye images were captured using Leica MZ10F microscope.

For wing mounting, adult wings were dissected from the hinge region of flies with the help of needle and scalpel and washed in isopropanol. The wings were then immediately mounted using wing mounting media. Wing images were taken using Brightfield Nikon Eclipse Ni microscope.

### Embryo Collection

Embryos were collected on a 2% agar plate supplemented with 0.2% propionic acid and yeast paste. Embryos were washed with distilled water, dechorionated in Sodium hypochloride, and fixed for 1 hr in 1:1 heptane and 4% paraformaldehyde solution. Embryos were then devitellinized by replacing heptane with methanol followed by vigorous shaking. Further washing was done in methanol. Devitellinized embryos were stored in methanol at −20°. Embryonic stages were identified as described in Campus-Ortega and Hartenstein (1985).

### Immunocytochemistry and confocal microscopy

*Drosophila* third instar larval tissues were dissected in cold 1X PBS (Phosphate Buffer Saline)and fixed in 4% paraformaldehyde for 20 minutes at room temperature. This was followed by four times washing of the discs using a washing solution (0.1% BSA in Tri-PBS), for 15 minutes each. Blocking was performed using a blocking solution (PBST with 8% Normal Goat Serum) for 30 minutes to 1 hour. Primary antibody was added and tissues were incubated with it for overnight at 4 °C. Next day, the tissues were again washed with washing solution 4 timesat room temperature for 20 minutes each. This was followed by blocking for 1 hour at room temperature. Tissues were then incubated in a secondary antibody for 90 minutes. After washing the tissues with the washing solution for four times and one wash in 1X PBS at roomtemperature, DAPI (1 µg/ml) was added for 20 mins to overnight in the dark. The tissues were finally dissected in 1X PBS and incubated in 1,4-diazabicyclo[2.2.2] for overnight. The next day, mounting of the discs was done and tissues were observed under LSM 780 laser scanning confocal microscope Zeiss [Carl Zeiss]. The images were processed in Adobe Photoshop 7. Mouse anti-Cut (1:100, 2B10), mouse anti-Wg (1:100, 4D4), mouse anti-Arm (1:100, N2 7A1), mouse anti-En (1:100, 4D9), mouse anti-Futsch (1:100, 22C10), mouse anti-Dl (1:100, C594.9B), mouse anti-NICD (1:300, C17.9C6), were procured from Developmental Studies Hybridoma bank, guinea pig anti-Sens (1:100) a kind gift from Prof. Hugo Bellen, Houston, USA. Department of Molecular and Human Genetics, Baylor college of Medicine; rabbit anti-Vg (1:100), rabbit anti DCP-1 (1:100, 9578) was procured from Cell Signaling Technology, rabbit anti-HA (1:100, SAB5600116) procured from Sigma-Aldrich were used as primary antibodies. Alexa Fluor 555 conjugated goat anti-mouse IgG (1:200), Alexa Fluor 488 conjugated goat anti-mouse IgG (1:200), Alexa Fluor 555 conjugated goat anti-rabbit IgG (1:200), Alexa Fluor 555 conjugated goat anti-rat IgG (1:200) and Alexa Fluor 488 conjugated goat anti-rabbit IgG (1:200) (all from Molecular Probes) were used to detect primary antibodies.

### RNA isolation and quantitation of transcripts

Total RNA was isolated from adult heads using TRIZOL reagent following the manufacturer’s recommended protocol (Sigma-Aldrich). RNA (2 μg) was treated with DNAse using 1.0 µl 10X reaction buffer, 0.5 µl DNAse (New England Biolabs) and 0.5 μl RNAse inhibitor (1,000 U/ml),and volume was adjusted to 10 µl using DEPC MilliQ water. The reaction mixture was incubated at 37°C for 1 h. For single-stranded cDNA preparation, 10 µl of DNAse-treated RNAwas mixed with 1 µl M-MuLV reverse transcriptase enzyme (New England Biolabs), 2 µl of 60 μM random primers, 2 µl of 10× M-MuLV buffer, 1 µl of 10mM dNTP and 1 µl of RNase inhibitor, and nuclease-free water was used to make the total volume to 20 µl. The mixture was incubated at 25°C for 5 min followed by a 42°C incubation for 1 hour. This cDNA was used asa template DNA for semiquantitative and RT quantitative (RT-qPCR). RT-qPCR was carried out as per the manufacturer’s protocol (Applied Biosystems). A total of 10 μl of the reaction volume included 5 μl 2× SYBR green, 0.25 μl of each forward and reverse primer, and 1 μl cDNA; PCR was performed using an ABI 7500 instrument. Data were normalized to rps17 before calculating the relative fold change.

Primers used for study are as follows:

mβ_RT_Fw 5’- ACCGCAAGGTGATGAAGC -3’

mβ_RT_Re 5’- CTTCATGTGCTCCACGGTC -3’

mδ_RT_Fw 5’- ATGGCCGTTCAGGGTCAG -3’

mδ_RT_Re 5’- CCATGGTGTCCACGATG -3’

mγ_RT_Fw 5’- GTCCGAGATGTCCAAGAC -3’

mγ_RT_Re 5’- GACTCCAAGGTGGCAACC -3’

m3_RT_Fw 5’- ATGGTCATGGAGATGTCC -3’

m3_RT_Re 5’- GCACTCCACCATCAGATC -3’

m5_RT_Fw 5’- ATGGCACCACAGAGCAAC -3’

m5_RT_Re 5’-TGTCCATTCGCAGGATGG -3’

m7_RT_Fw 5’- GGCCACCAAATACGAGATG -3’ m7_RT_Re 5’- CAT CGC CAG TCT GAG CAA -3’m8_RT_Fw 5’- GGAATACACCACCAAGACC -3’ m8_RT_Re 5’- CGCTGACTCGAGCATCTC -3’ rps_RT_Fw 5’-AAGCGCATCTGCGAGGAG-3’ rps_RT_Re 5’-CCTCCTCCTGCAACTTGATG-3’ Dl_RT_ FP 5’- CATCGTGCAGGTTCACAGTT-3’ Dl_RT_Re- 5’ GTGGCCTGGTAGTGCTTTAG-3’

### Climbing assay

A cohort of n > 15 age-matched adult female flies for each genotype was collected and transferred to a graduated cylinder with a height of 20 cm and a diameter of 2 cm. The cylinder was tapped to collect the flies at the bottom, and the movements of the flies were recorded for a duration of 10 s. These procedures were repeated at least 10 times.

### Acridine Orange staining

Acridine Orange was used to detect apoptosis in developing *Drosophila* imaginal discs. Imaginal discs from third instar larvae of the F1 progeny from different crosses were dissected out and washed briefly in cold 1X-PBS (pH 7.2). These discs were incubated in Acridine Orange (1 µg/ml; Sigma) for 1 min, followed by two gentle washes in 1X-PBS. Then the discs were mounted and observed immediately under Nikon Eclipse 80i fluorescence microscope ata 488 nm emission filter.

### Statistical analysis

Intensity profiling in the *Drosophila* imaginal discs and quantification of the adult wing area was done using Image J. 3-5 imaginal discs were used for the quantification purpose in each case. Integrated density/area of the domain indicated the intensity of the staining in confocal images. The RGB plot profiles feature of Image J was also used to measure the intensity of Notch signaling components in different genetic combinations compared to the internal control.

RGB plot profiles feature from plugins section was then used to plot intensity graphs. The errorbars in the graphs denoted the standard error of the mean value from the replicated experiments. Each dataset was repeated at least three times. T-test and Oneway analysis of variance (ANOVA) with Tukey’s multiple comparison post-test were employed to determine the significance of the level of difference among the different genotypes. p-value < 0.05 was accepted as statistically significant.

## ACKNOWLEDGMENTS

The authors extend sincere thanks to Prof. Spyros Artavanis-Tsakonas (Harvard Medical School), Prof. Ella Preger-Ben Noon (Janelia Research Campus, Ashburn, VA, USA), Prof. Hiroshi Shibuya (Simon Fraser University, Canada), Prof. Alexey Veraksa, (University of Massachusetts Boston, USA) and The Bloomington Stock Center for providing fly stocks. We heartfully acknowledge our gratitude to Prof. Hugo Bellen, (Department of Molecular and Human Genetics, Baylor college of Medicine, Houston, USA) and Developmental Studies Hybridoma Bank, University of Iowa, USA for stocks for antibody support. We acknowledge Dr. Nalini Sachan for *UAS-HA-Chip* fly generation. We also acknowledge the confocal facility of DBT-BHU-ISLS, and CDC, Banaras Hindu University.

## AUTHOR CONTRIBUTIONS

AM and JS designed the experiments. JS, DV, and BS did all the experiments. AM, MM, MSPand JS analyzed all the data. AM and JS wrote the manuscript.

## DATA AVAILABILITY

The datasets supporting the conclusions of the article are included within the article.Dataavailable upon request.

---

**Supplementary Figure 1. Awh regulated Wingless signaling is mediated through Notch signaling.** Overexpression of Awh in AP boundary shows reduced Wg (A2, High magnification in A1) GFP marks the domain of Awh overexpression (A3) DAPI shows the defect in wing morphology (A4) Merged image is shown (A5). Overexpression of NICD in AP boundary shows enhanced expression of Wg (B2, High magnification in B1) GFP marks the domain of Awh overexpression (B3) DAPI shows the wing morphology (B4) Merged image is shown (B5). Co-expression of Awh and NICD in AP boundary rescue Awh mediated Wg downregulation (C2, High magnification in C1) GFP marks the domain of Awh overexpression (C3) DAPI shows the rescue in Awh mediated disruption in wing disc morphology (C4) Merged image is shown (C5). Graph shows the intensity of Wingless in different combination (D).

**Supplementary Figure 2. Spatial regulation of Awh.** Awh overexpression showed reduced expression of Notch target Cut and Wg at DV boundary, ectopic Cut and Wg was observed in dorsal compartment, no change is observed in ventral region (A1,B1) GFP marks the Awh gain-of-function clone (A2,B2) Merged images are shown, blue is DAPI (A3, B3).

**Supplementary Figure 3. Chip rescue Awh mediated phenotype.** Overexpression of Awh using *vg-GAL4* and *ptc-GAL4* leads to severe wing disc deformities such as small wing disc (A1) and duplication (B1). The phenotypes were significantly rescued upon bringing Chip overexpression line in background (A2, B2).

**Supplementary Figure 4. Overexpression of Awh led to cell death.** Overexpression of Awh using *en-GAL4, dpp-GAL4, ptc-GAL4, ap-Gal4, vg-GAL4* led to increased dying cells marked with Acridine Orange.

**Supplementary Figure 5. Overexpression of Awh led to Caspase mediated cell death.** Overexpression of Awh in ventral region leads to significant cell death as shown with dcp-1 expression (A1). Downregulation of *Drosophila* Caspase, Dronc and overexpression of inhibitor caspase p35 in Awh overexpression background rescue the cell death (B1, C1). GFP marks the domain of Awh overexpression (A2,B2,C2) and merged images shows expression of dcp-1 and respective domain (A3,B3,C3).

**Supplementary Figure 6. Inhibiting Caspase mediated cell death can not rescue Awh mediated wing margin defects.** Overexpression of Awh leads to severe wing defects (A), inhibiting cell death using UAS-p35 in the background rescue the wing size but not the wing margin (B).

