## Supplementary figures and images for "Notch and LIM-homeodomain protein Arrowhead regulate each other in a feedback mechanism to play a role in wing and neuronal development in *Drosophila*"

### Supplementary Figure. S1

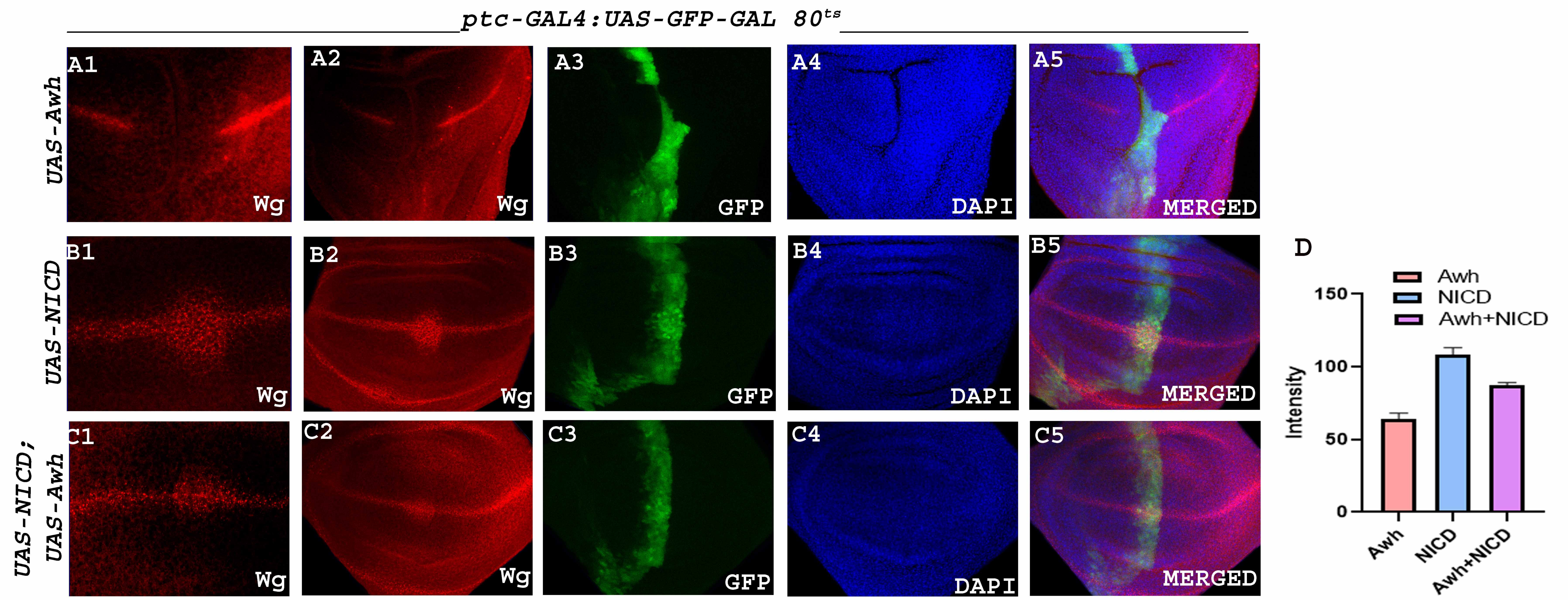

### Supplementary Figure. S2

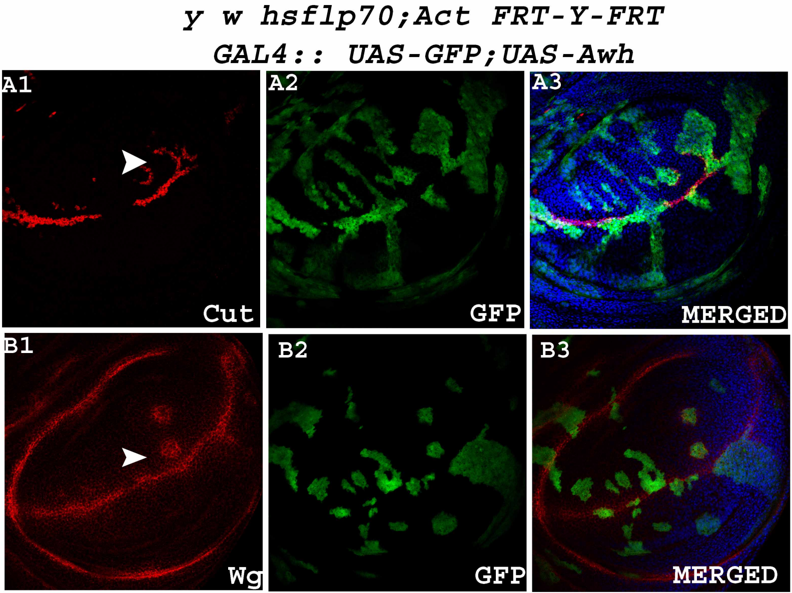

### Supplementary Figure. S3

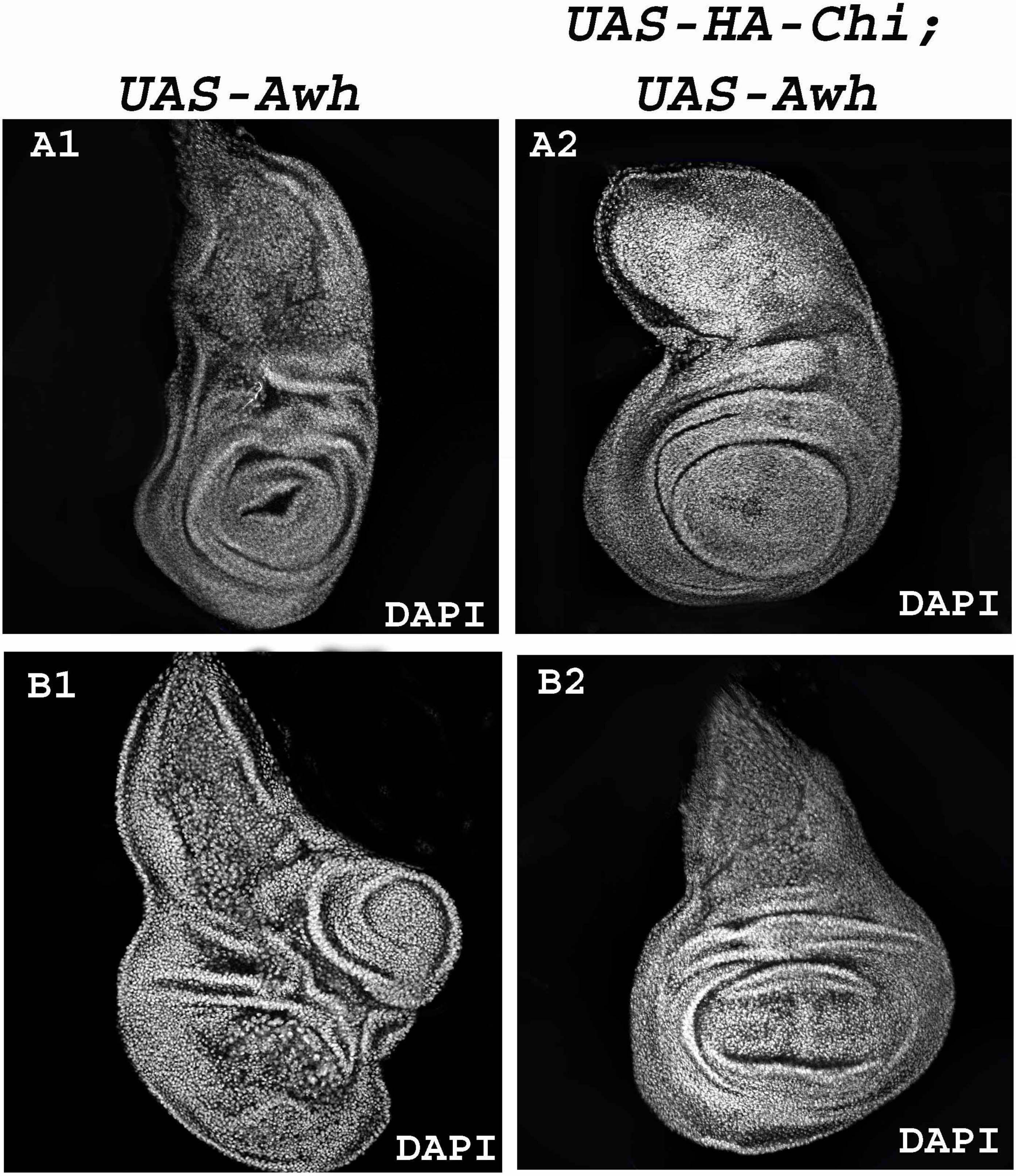

### Supplementary Figure. S4

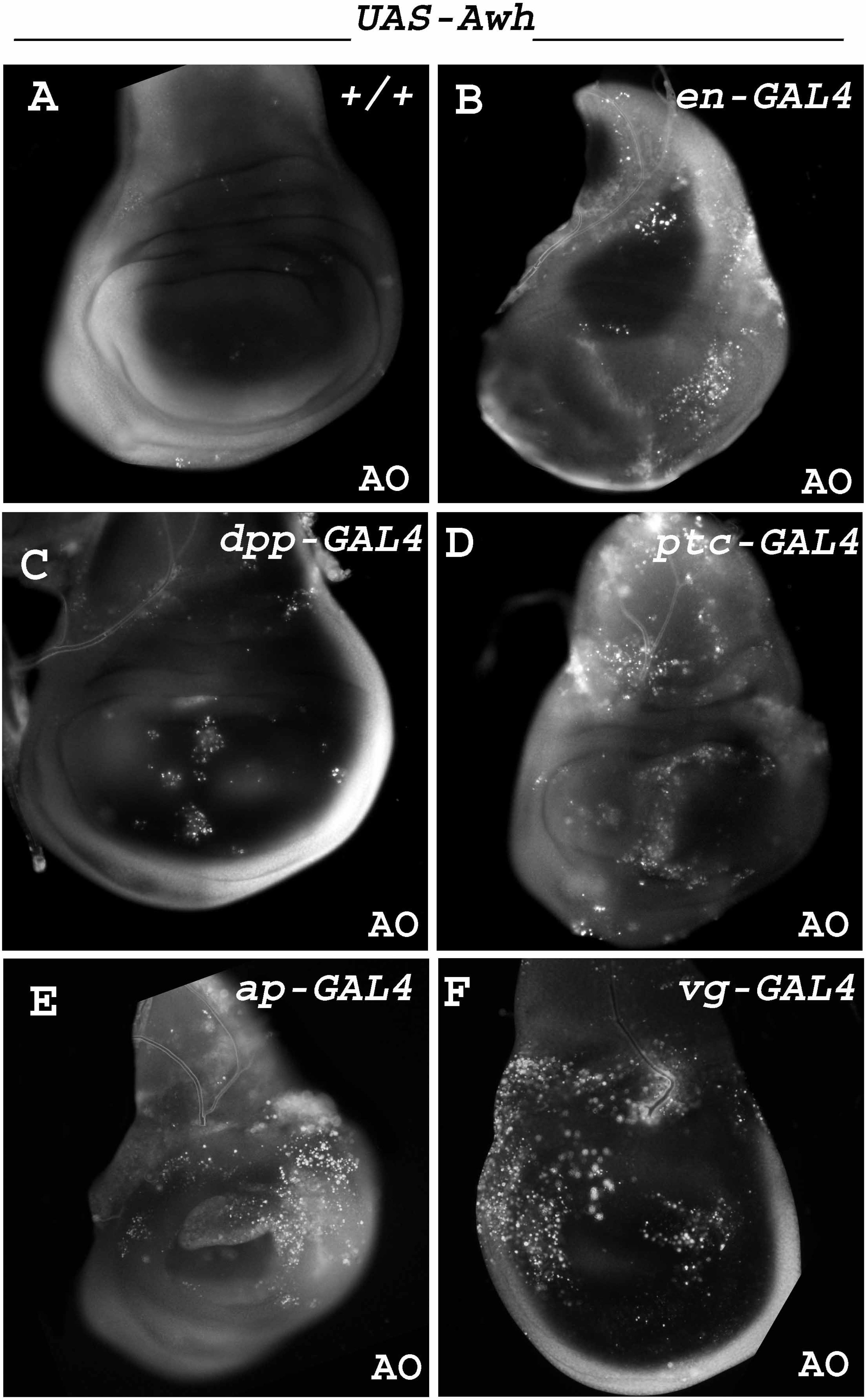

### Supplementary Figure. S5

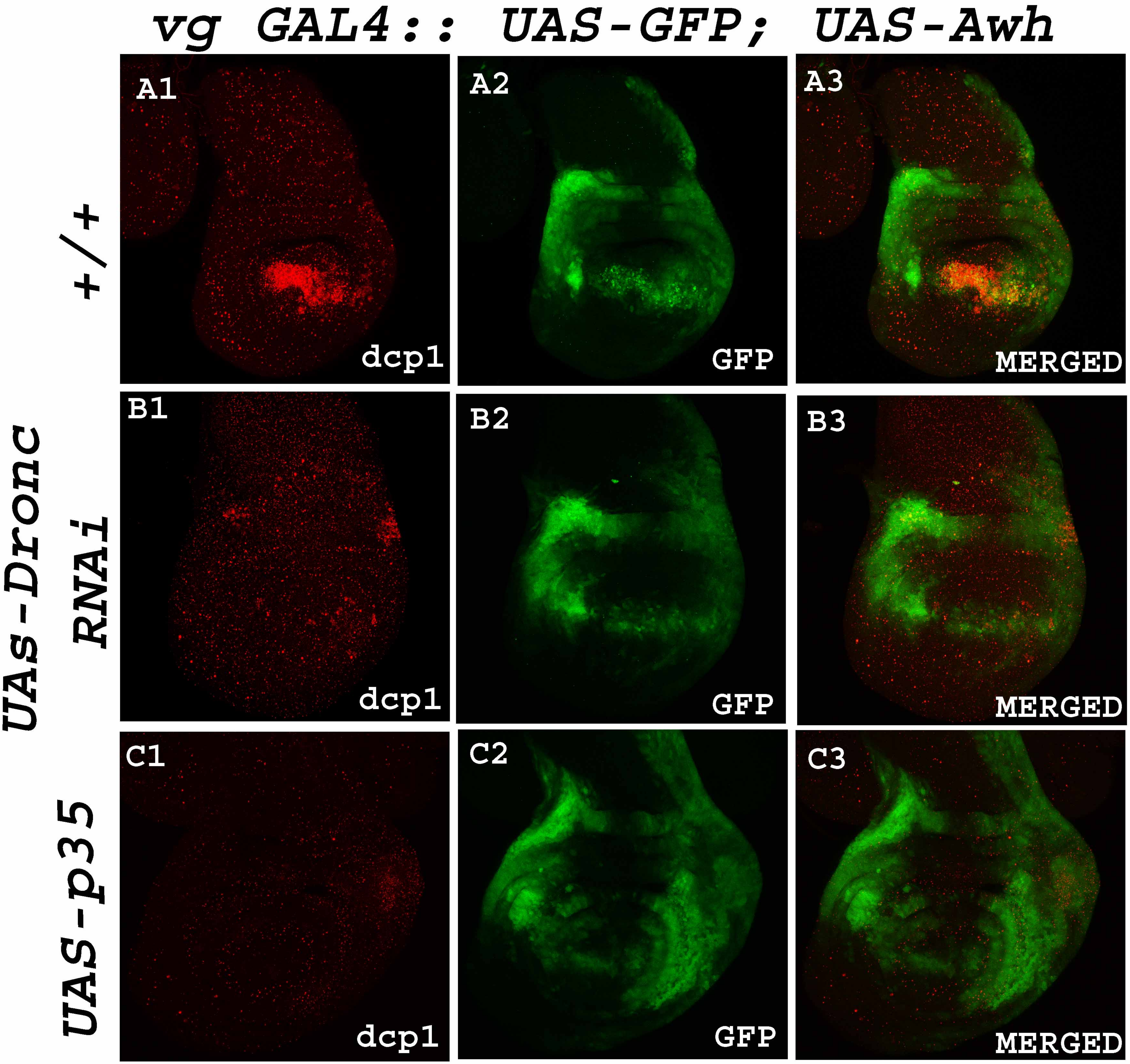

### Supplementary Figure. S6

*vg GAL4; UAS-Awh* | *vg GAL4; UAS-Awh/ UAS-p35*

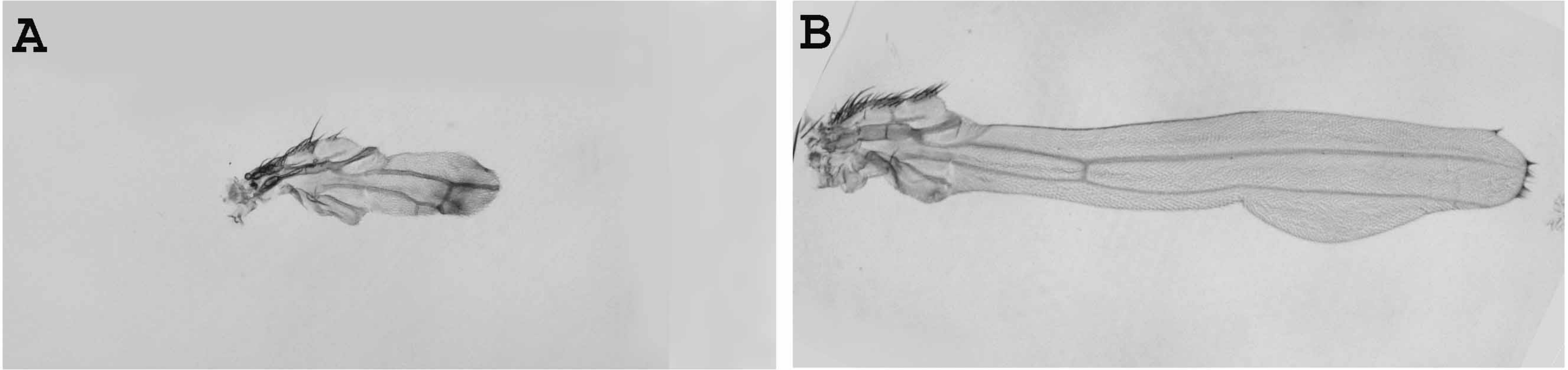
